# 1000 years of population, warfare, and climate change in pre-Columbian societies of the Central Andes

**DOI:** 10.1101/2022.11.24.517643

**Authors:** Mauricio Lima, Eugenia M. Gayó, Andone Gurruchaga, Sergio A. Estay, Calogero M. Santoro

## Abstract

Different Andean societies underwent processes of expansion and collapse during propitious or adverse climate conditions, resource boost or depletion along with population variations. Previous studies have emphasized that demographic collapses of polities in the Central Andes Area were triggered by warfare and the negative impacts of fluctuating climate (droughts) on crop productivity. Nevertheless, the interactions between climatic variability, demography and warfare have been less thoroughly evaluated. We develop population dynamic models to test feedback relationships between population growth, climate change and warfare in the Central Andes, where considerable regional hydroclimate variations have occurred over a millennium. Through population models, we found out that the rise and demise of social polities in the northern coast of the Central Andes appear to be a consequence of climate change. In contrast, for the highlands of Peru and the Titicaca basin, population models suggest that warfare intensity has a negative effect on population growth rates.

## 1 Introduction

Holocene climate variability affected the trajectory of ancient civilizations across the globe within complex processes of mutual influence [1], where human societies play key role in transforming historical landscape, resulting in enhancing richness, soil fertility, equitability biodiversity of nature and landform heterogeneity [2, 3].

Different complex societies after developing several action to transform their agrarian, agro-pastoral or agro-marine economies landscape, underwent collapse during periods of adverse climate conditions, resource depletion and high population levels [4-6], which could not be overthrow by human technological and social actions. The vulnerability of these economies could increase due to the “threat multiplier” effect of changes in the frequency and duration of extreme climate events in straining social instability in states with weak governance [7-9] and limited technological solutions. However, such causal links between climate change, population growth, critical resources and social conflicts are still debated [10-12]. Causality analyses suggest that past demographic collapses in China and Europe were triggered by social unrest, which was ultimately modulated by the negative impacts of prolonged cooling events on crop productivity and per capita food supply [13, 14]. Conversely, nonlinear dynamical models attest to a much more active role of demography in driving warfare and population collapses via offer-demand dynamics over strategic resources [15-17]. That is, population demises could arise from the endogenous interplay between population size/pressure and social conflict. Nevertheless, to the best of our knowledge, the interference of changes in climate conditions in nonlinear dynamics between demography and social conflict has not been empirically evaluated.

Here, we develop a quantitative approach to test long-term feedback relationships between population growth, climate change and warfare by implementing population models based on the principles of nonlinear dynamical systems. We focused on the Central Andes area (Fig. 1A), in which several state level polities have risen and collapsed under diversified ecogeographic zones, from the cold semiarid Andean highlands (the southern centers area) to the extreme hyperarid Pacific coast of South America (the northern centers area), where people developed transformative complex productive technological systems. Parallelly, the region has experienced considerable variations in regional hydroclimate conditions over the last millennium [18-21]. Thus, much attention has been given to the potential role that changes in water availability and technological management could have played in the rise and fall of Andean state level polities across the rather arid landscape that characterizes the western Andean slope. For example, the demise of the Tiwanaku and Wari empires in AD ∼1000 has been linked to an intense regional megadrought that limited crop production over the Andean highlands [19, 22-24]. The collapse of the State Moche societies in approximately AD 900 [25] has been attributed to the loss of arable lands caused by repeated drought or flood cycles and sand dune invasion due to severe of El Niño events that affected their territories in the northern Peruvian coast, which is usually dominated by arid conditions [26-29]. Although most of available evidence indicates that these demises were accompanied by increased warfare intensity [30-34], the interaction of internal conflicts has been poorly considered. Still, McCool et al. [35] have recently explored the relationship between demography, hydroclimate and conflict in groups that dwelled mid-elevation valleys from the southern centers area at 700-1400 AD. Based on statistical causality analyses, these authors found that chronic conflict in the Nasca region was indirectly linked to climate, but directly to increased population size/pressure, which was ultimately set by the positive effect of wetter conditions on the food production in an arid region with limited availability of croplands.

**Figure 1:**
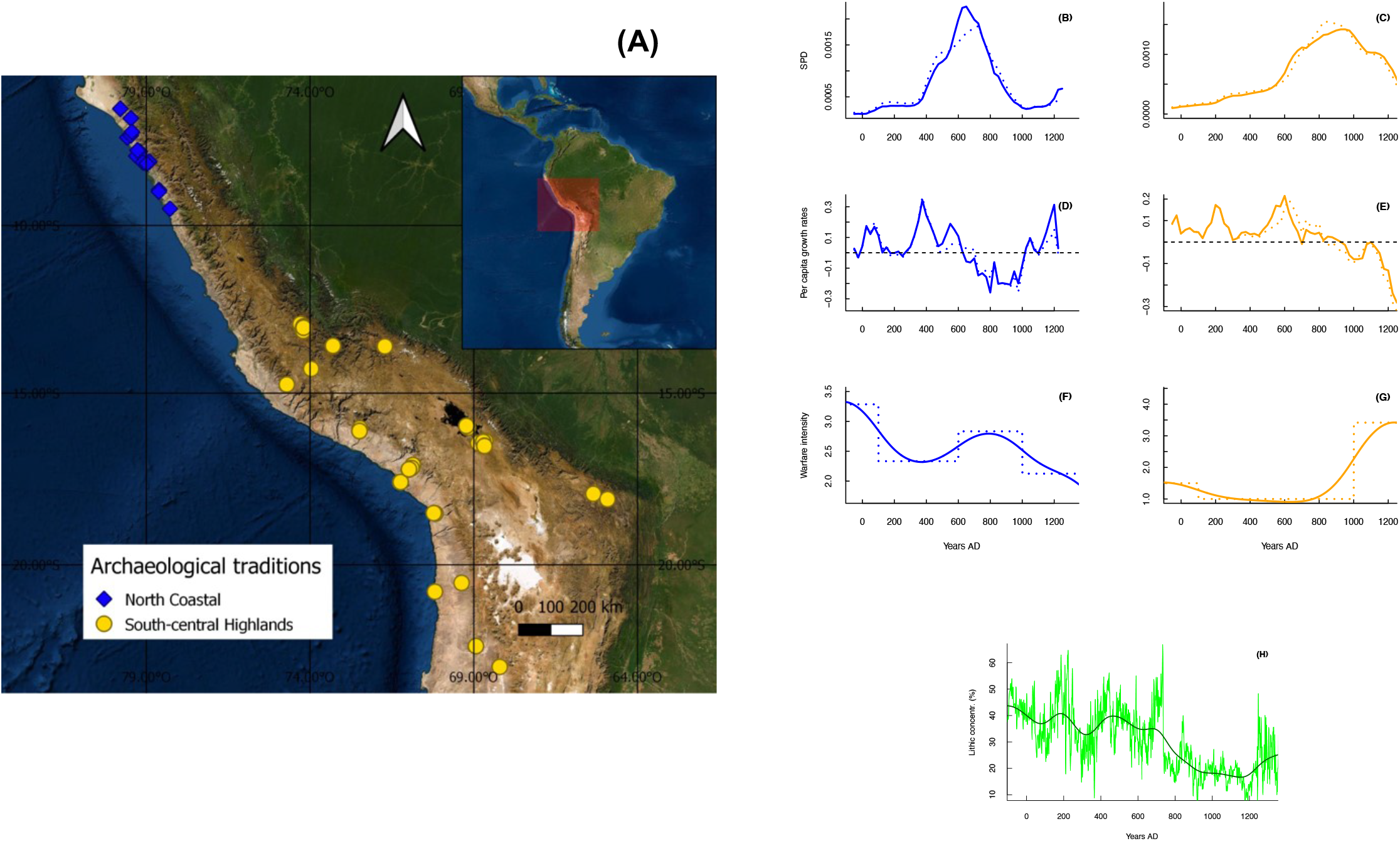
Central Andes archeological sites and the analyzed time series. (A) - Map showing the locations of archaeological sites providing radiocarbon data for each cultural tradition. (B) and (C) - Normalized (solid curves) and unnormalized (dashed curves) SPDs for northern (blue curve) and southern (orange curve) sociocultural areas. Vertical dashed bars indicate declining population growth rates. Note that colors of curves that describe SPDs for each tradition are maintained in panels below. (D) and (E) - Calculated growth rates based on normalized (solid curve) and unnormalized (dashed curve) SPDs. (F) and (G) - Raw (dashed curves) and smoothed (solid curves) time series for warfare intensity in each sociocultural area. (H) - Proxy time series for regional hydroclimate conditions. The green curve shows annually resolved lithic concentrations (see [21]), and the solid dark curve denotes the corresponding smoothed time series.

We fitted population dynamic models to empirically capture the interplay between long-term population size/pressure, warfare intensity and regional hydroclimate conditions (see Methods). To account for past demographic levels over the last 2000 years, we used as proxy data of Summed Probability Distributions (SPDs) for archeological radiocarbon dates aggregated in a novel chronometric database for the region. Our models were supplemented with time series for internal conflicts based on the dataset for defensive settlements developed by Arkush and Tung [30]. To explore the effect of anomalies in hydroclimate conditions, we considered a high-resolution proxy for long-term changes in the activity of the El Niño Southern Oscillation (ENSO), which is the main driver for hydroclimate, frequency of extreme weather events, bioproductivity and agricultural production over the western Andean slope at the interannual scale [36-39]. Linguistic, genetic and archeological evidence attests to two relatively independent centers of cultural and sociopolitical development across the Central Andes area that encompasses from the modern Ecuadorian-Peruvian border to the Lake Titicaca basin in Bolivia [40, 41]. The northern center is represented by agro-marine social structures that dwelled on the northern Peruvian coast (hereafter the northern sociocultural area; Fig. 1A). The southern nuclei include agro-pastoral societies from the highlands of southern Peru and the Titicaca Basin (i.e., the southern sociocultural center, Fig. 1). Consistent with such social structures, we evaluated dynamical interplays independently for the northern and southern centers.

## 2 Methods

### 2.1 Paleodemographic data

We generated a database of 382 radiocarbon dates of terrestrial samples (i.e., charcoal, plant remains, and wood) collected from 70 archaeological sites along the Central Andes Culture Area (Fig. 1A). We followed criteria proposed by Bird et al. [42] to assemble archeological ^14^C dates into a database that provides information on long-term paleodemographic trends in a given area. The northern sociocultural area is represented by 125 ^14^C dates from 13 archeological sites unequivocally assigned to the Moche tradition. In contrast, 257 dates from the Wari and Tiwanaku archaeological sites (n=57) serve as the southern sociocultural center. The resulting dataset covers the period of 50 to 1300 AD, concatenating existing repositories and previously published data (e.g., [43-45]. The dataset is available through the GitHub account of the PEOPLE 3000 Working Group of PAGES (https://github.com/people3k) and is thus indexed as an update of the P3K14C dataset [42]. Sampling intensities are 0.14 and 0.26 dates/100 yr/100 km^2^ for northern coast and south-central highland traditions, respectively. These intensities are within the range reported in other works that examine paleodemographic trends and patterns for different regions of South America (0.13-0.22 dates/100 yr/100 km^2^; [46].

To estimate demographic levels over time, we calculated independent SPDs of calibrated radiocarbon dates for each sociocultural area. Theoretical and empirical studies indicate that SPDs represent a good proxy for relative changes in population growth and human energy consumption [47]. Radiocarbon data were processed in the Rcarbon package for R [48]. All data were calibrated using the SHCAL20 calibration curve [49]. We controlled the edge effect by calibrating and summing dates over the time range of 50 to 1500 AD. To offset the contribution of intra-site overrepresentation and calibration biases, we applied a bin size of 50 years and a 100-year rolling mean. Following recommendations made by [48], we generated SPDs based on normalized and unnormalized radiocarbon dates. Logistic null models on normalized and unnormalized SPDs were run to test positive or negative deviations from simulated envelopes via “calsample” and “uncalsample” methods, respectively [48]. Before the implementation of population dynamic models, normalized and unnormalized SPD time series were sectioned into time-step intervals of 25 years. This procedure allowed us to capture only major population trends, avoiding a high-frequency noise source of variability [5].

### 2.2 Warfare data

Original data of warfare intensity come from the review by [30], which collates archeological evidence identifying threats or harms that past Central Andes populations suffered in times of conflict (Fig. 1D). The review by [30] considers frequencies of violent skeletal trauma and defensive settlements. We selected data for defensive settlement because the chronological and spatial resolution for this proxy is well resolved in our studied areas. In practice, the frequency of defensive settlements represents the actions taken by a given group to face the threat of attacks from other populations. Under the classification of [30], a defensive settlement is understood as a settlement localized in areas that are difficult to access because of its geographic characteristic (e.g., hilltops) that may or may not have constructive fortifications (structural defenses or fenced villages), or other strategic architectural devices (shelters, walls, ditches, checkpoints), complementary to enhance its unassailability or to reduce their vulnerability. The basic data used for the frequency of defensive settlement were obtained by coding numerically whether these were absent (0), present (1), or common (2) during a given period over each area. Hence, frequencies within this raw time series range from 0 to 6. Because warfare data are grouped into broad chrono cultural periods of the Central Andes area (e.g, Formative, Early Horizon, Early Intermediate Period, Middle Horizon, Late Intermediate Period, Late Period), we translated these discrete and large time step data into an annual time index. Then, the resulting time series was smoothed using a cubic smoothing spline function [50] with a spar (smoothing) parameter of 0.90. We then resampled the data at 25-year intervals to match the SPD time series (Fig. 1E).

### 2.3 Hydroclimate data

Even when there are different hydroclimate reconstructions for local hydroclimate over restricted areas of the western Andean slope, our perspective is based on the premise that proxies for overall drivers best describe large-scale conditions (e.g., “packages of weather” *sensu* [51]). This approach relies on the fact that many available reconstructions for local hydroclimate are temporally discontinuous and/or poorly constrained due to insufficiently resolved paleoclimate records. That is, we assume that reconstructions for past changes in ENSO activity serve as proxies for long-term hydroclimate conditions over the Central Andes area. The selected ENSO proxy is the annually resolved concentration of terrestrial clasts in a marine core (SO147-106KL) retrieved ∼80 km off the Peruvian coast (12°S, [21]). We used lithic concentrations instead of photosynthetic pigments or alkenone records, as riverine sedimentary flushing events into the Peruvian shelf reflect variations in regional hydroclimate conditions (Fig. 1E). Specifically, lithic concentrations in this sediment core depend on the intensity/magnitude of continental precipitation. Thus, decreased/increased clast accumulations in the marine core are associated with prevailing La Niña/El Niño conditions over the Tropical Pacific and, in turn, negative/positive rainfall anomalies and a reduced/increased frequency/intensity of flooding events along the northern Peruvian coast [21]. Reduced/increased lithic concentrations, and in turn prevailing La Niña-like/El Niño-like conditions, respectively are associated with increased rainfall over the Andean highlands.

Raw data for clast percentages in the SO147-106KL core are presented at an annual timescale (green curve in Fig. 1E), thus capturing high-frequency climate variations. Because we are interested in the long-term signal for mean hydroclimate conditions (i.e., wet interludes versus droughts), we smoothed the raw time series by fitting a cubic spline function with a spar parameter of 0.65 (dark curve in Fig. 1E). The smoothed time series was resampled at 25-year intervals to match the SPD time series.

### 2.4 Population dynamic models

We applied a modeling framework that assumes a coupled dynamical system between two endogenous variables and an exogenous variable. The endogenous variables are population dynamics (N_t_) and warfare intensity (W_t_) [15, 17]. The exogenous variable is represented by long-term climatic variability (C_t_). We assume that changes in hydroclimate conditions will affect long-term food production or crop yields, in turn modifying the equilibrium density (*k*, also called carrying capacit*y*) of a given society. Thus, *k* is set by the available amount of arable land for farming and herding and concomitant technologies (yield per unit of area). As the population approaches *k*, all available resources will be used (e.g., cultivable land, water management). Further increases in population size will immediately result in lower average consumption rates. However, these societies may face long-term changes in climate conditions that modulate resource availability (i.e., the amount of arable land or yield per unit of area). The explanation relies on the combination of Malthusian theory [52] with climatic variability as an exogenous forcing factor. Climatic variability determines agricultural land carrying capacity, which affects the population growth of societies [4, 5]. Lateral perturbations result from exogenous factors (e.g., climate) that act on population equilibrium *k* and cause nonadditive effects. In the absence of warfare, the dynamic of this system can be defined as a logistic equation with lateral perturbation effects [53]:

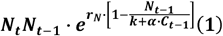

*N*_*t*_ denotes the human population size at time *t*; *r*_*N*_ is a positive constant representing the maximum per capita reproductive rate, *k* is the “carrying capacity” of the system, α is the climatic coefficient and C_t-1_ is the climatic variable or the proxy influencing food production. In fact, the ratio N/resource availability value in Equation 1 is the proxy of “population pressure” defined as the relationship between population size relative to available resources [54-56]. However, the rise in population in complex societies might increase the frequency of warfare and conflicts, creating a mutual feedback structure among population growth and warfare intensity [e.g., 17]. Therefore, the population dynamic model of Equation 1 can be modified to include a term for warfare intensity or frequency:

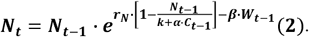

where α is the climatic effect on crops or land productivity, and *β* is direct mortality caused by warfare. The model can be modified to introduce the effects of both climate change and warfare intensity on land productivity:

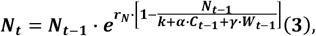

Parameter γ represents the negative effects of warfare intensity on land productivity [17]. We estimate that two processes could drive the dynamic of warfare intensity in complex societies. First, large population sizes increase opportunities for encounters between social groups, in turn placing demographic pressures on land use and productivity. Second, an increased probability of armed conflicts between social groups could also arise from reductions in land productivity or area due to the occurrence of adverse climate conditions such as prolonged droughts or more frequent floods. The impact of these extreme hydroclimate events is especially important along the arid western slope of the central Andes, as they are capable of triggering yield losses in several traditional crops [37]. Therefore, to account for the dynamics of warfare intensity (W), the starting rate of conflict *λ* is assumed to be proportional to population density, and by appealing to the mass-action law, there is an exponential rate of conflict decay μ [e.g., 17]. The basic model for warfare intensity dynamics is as follows:

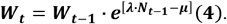

The impact of climate fluctuations was incorporated into this model by assuming that harsh hydroclimate conditions result in reduced crop productivity and increased warfare intensity. There are two different hypotheses; one proposes a simple additive effect of climate.

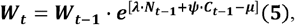

where constant Ψ represents the negative/positive effects of climatic change on warfare dynamics. On the other hand, as in Equation 1, we can assume that the ratio population/resource availability is the proxy of “population pressure” defined as the relationship between population size relative to available resources [55]. Hence, the dynamics of warfare might be driven by the ratio of population size/land productivity.

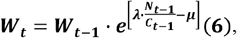

In sum, we expect the population rate of change (log N_t_ – log N_t-1_) to be negatively affected by warfare, while population size will positively influence the warfare rate of change (log W_t_ – log W_t-1_). Hydroclimate conditions are expected to have an exogenous perturbation effect on this coupled relationship.

### 2.5 Statistical Analyses: *Nonlinear regression analyses and model comparison and validation*

The parameters of Equations 1-6 were estimated through nonlinear least-squares fitting procedures using the nls (nonlinear least squares) library on the R platform. To fit these nonlinear models, we used the population (r_t_) and warfare (w_t_) rates of change as response variables by log transforming Equations 1-6.

Because we use smoothed time series for SPDs (population levels), warfare intensity and hydroclimate conditions, a temporal autocorrelation is included, which creates some problems when using standard statistical tools for assessing the goodness of fit of the models. To compare the statistical models, we used different and complementary approaches. First, we selected the models by measuring the Bayesian Akaike information criterion (BIC) [57], and models with the lowest BIC values were selected. Second, we compare and validate the models by simulating the total trajectory predictions initiated with the first observed value of the time series and running the algorithm using each model with their estimated parameters to obtain the time series’ remaining simulated values.

In all simulations, the accuracy of predictions was assessed using coefficient of prediction σ^2^ [58]

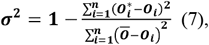

O_i_ is the observed data from the testing dataset, O_i_^*^ denotes the model predictions, Ō is the mean of the observations, and n is the number of data to be predicted. Coefficient of prediction σ^2^ is 1 when the predicted data are equal to the observed data, 0 when the regression model predicts and the data average, and negative if the predictions of the model are worse than the data mean.

## 3 Results

The raw SPDs for normalized and unnormalized radiocarbon dates from both sociocultural areas are practically identical, thus reproducing equivalent overall patterns for past population levels over time (Fig. 1B). The magnitude and amplitude for peaks (population increased) and troughs (population declines) in raw SPD curves vary slightly when normalized and unnormalized ^14^C-dates are summed. The same is true for the extent and duration of positive and negative significant deviations from logistic null models (Fig. S1).

We document striking temporal variation in population dynamics for the period of AD ∼ 50-1300 in both the northern and southern center areas (Fig. 1 B-C). The northern and southern centers experienced demographic levels peaking at AD ∼650 and AD ∼1000, respectively (Fig. 1 B-C). This implies that these sociocultural areas underwent sustained phases of population growth that lasted between 400 and 500 years (Fig. 1 D-E). Analyses of temporal changes are consistent with this demographic pattern, suggesting rapid increases in regional population levels from AD ∼400 to AD 550 (Fig. 1D - E), during the so-called Middle Horizon period (AD ∼600-1000). This overall growth phase matches with the rise of Wari, Tiwanaku and Moche state level polities with their characteristic iconographic styles [43-45].

For the northern centers area, there was a rapid growth transition starting at AD ∼300-350 and then a demographic collapse occurring at approximately AD 900 (Fig. 1D). The latter coincides closely with the estimated chronology for the end of the material culture that characterized the Moche tradition (26). In the southern centers area, a sudden increase in growth rates took place by AD ∼500-600. This phase was followed by strong population decline centered at AD ∼1100–1200 (Fig. 1E), which appears related with the timing of the fall of the Tiwanaku and Wari traditions [43-45], and strong climatic deterioration [19, 24, 26].

Corresponding data for northern and southern centers are constrained to AD 50-925 and AD 250-1225, respectively (Figure S2 and S3 B-D). Hence, we can evaluate whether distinct boom-and-bust dynamics that characterized different past Andean societies (e.g., [6]) were also present in the Central Andes Area. We verify that this procedure captures the core processes of such a dynamic and that these cultural areas indeed endured socio ecological rise and fall from AD ∼50-1250 (Figure 1 A-D).

At first glance, population dynamics at each of the sociocultural centers area follow long-term variations in warfare intensity and hydroclimate (Fig. 1F-H). Positive population growth rates prevailed over the northern centers (AD 400-600; Fig. 1D) when warfare intensity was relatively low (Fig. 1F), and clast concentrations in the SO147-106KL core were high (>35%, Fig. 1H). The peak in population growth rates (AD 400-500) coincides with a period of lowest conflict intensity and increased coastal rainfall. In contrast, population decline is observed from AD 500 to 600 as warfare intensity steadily increases, as interpreted by the proliferation of cranial trauma among adults [30, 31]. This also occurred during a period in which the Eastern Pacific went into El Niño-like conditions (AD 400-500) but characterized by an incipient decline in rainfall and flood activity (Fig. 1H). From AD 600 to 900, population decline occurred (Fig. 1 B and D), warfare intensity continued to increase (Fig. 1F), and the overall trend in lithic concentrations suggests reduced coastal precipitation (Fig. 1H).

The warfare intensity over the southern sociocultural centers area shows an increasing trend (Fig. 1G). The lowest values occurred in AD 600, concomitant with a period of population increase (Fig. 1E) and a long-term tendency toward a La Niña-like conditions (Fig. 1I), leading to rainfall increase over the Andean highlands. Around approximately AD 900, population began to decline, and conflict intensity increased rapidly. Such coupling between warfare and population dynamics, however, occurs despite the Eastern Pacific remaining locked into La Niña-like wet conditions (Fig. 1I).

Population dynamic models (Eqs. 1-6) highlight important differences between both northern and southern sociocultural centers areas on the long-term demographic/hydroclimate/conflict interplay. For instance, best-fitted population models either in BIC or R^2^ values (Table S1) for the northern sociocultural centers include population sizes and variations in hydroclimate. Total trajectory predictions from a model that includes climate as a lateral perturbation effect are considerably better than predictions from models that consider warfare effects (Figure 2 A-D, Table S1). Moreover, warfare dynamics are mainly determined by the additive combined positive effects of population size and hydroclimate (lower BIC and higher R^2^ values) (Table S1). Changes in the intensity of warfare are loosely related to population increase over the northern sociocultural center areas (Table S1).

**Figure 2:**
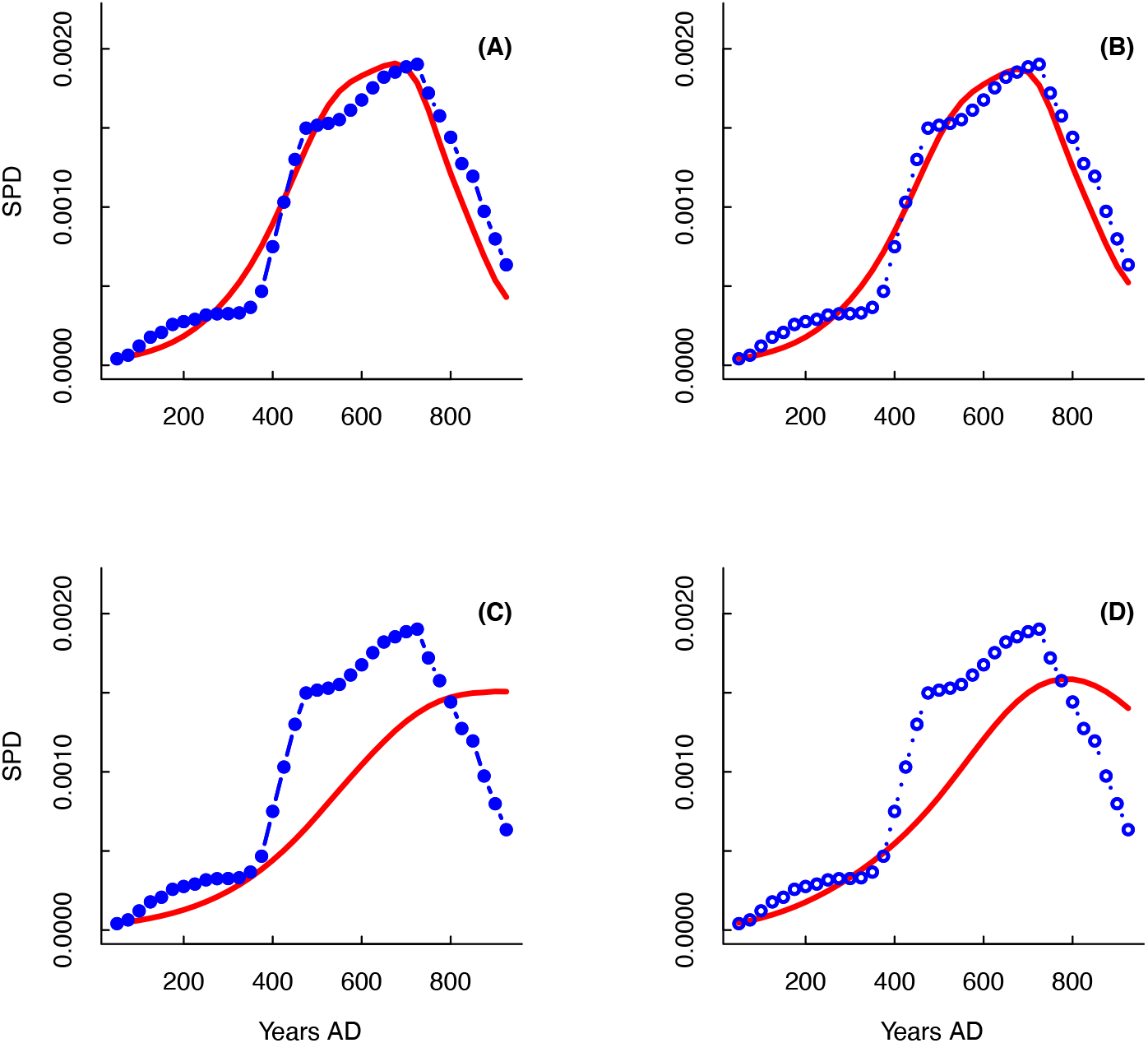
Predicted and observed population dynamic at Northern sociocultural area. Comparisons between population dynamic model-predicted trajectories (red solid curve) and observed SPD data (normalized SPD: solid line and blue dots; nonnormalized SPD: dotted lines and blue dots) for the northern sociocultural area. Panels (A) and (B) show model predictions that include hydroclimatic variation for normalized and unnormalized data, respectively. (C) and (D) show predictions for models including warfare effects for normalized and unnormalized data. Further details on the model predictions are provided in Table S2.

In the case of the southern sociocultural centers area, demographic dynamic models fitted to SPD data clearly demonstrate that warfare intensity has a negative effect on population growth rates. Those models that incorporate conflict explain more than three times the variance in population growth rates than models including hydroclimate conditions as predictors (Table S2). In fact, total trajectory predictions indicate that warfare is the best predictor and main driver of population dynamics over these sociocultural centers (Figure 3 A-D, Table S2). Likewise, warfare dynamics are mainly determined by the positive effect of population sizes (Table S2). Models that only consider demographic levels (i.e., SPD data) as explanatory variables explain over 80% of the change rate for conflict intensity (Table S2). Nevertheless, models including the effects of population pressure display lower BIC and higher R^2^ values, indicating that the ratio of population/water availability is the most important factor behind the warfare dynamic (Table S2). Thus, our results attest to the reciprocal influences of population and warfare in the southern sociocultural centers area where warfare has a negative effect on population growth rates, while population positively affects warfare growth rates.

**Figure 3:**
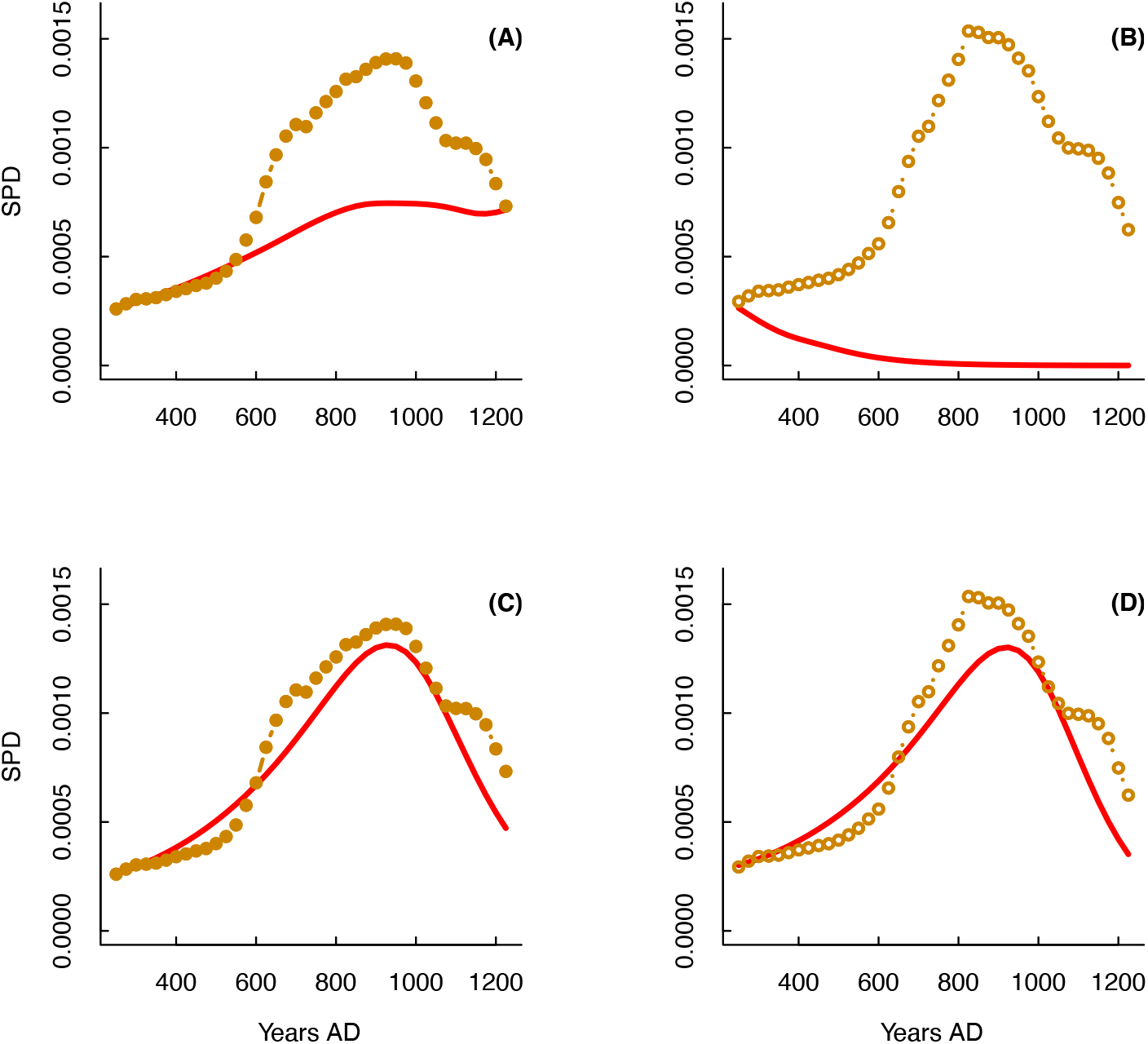
Predicted and observed population dynamic at Southern sociocultural area. Comparisons between population dynamic model-predicted trajectories (dark dashed curve) and observed SPD data (normalized SPD: solid curves; nonnormalized SPD: dotted orange curves) for the southern sociocultural area. Model predictions for the effect of changes in hydroclimate are shown for normalized and unnormalized data in Panels (A) and (B), respectively. Predictions of models that include warfare effects for normalized and unnormalized data are shown in Panels (C) and (D). Gray shaded areas show 95% confidence intervals for model predictions. Further details on the model predictions are provided in Table S3.

## 4 Discussion

Our results derived from a population dynamic approach suggest that the rise and fall of complex societies in the Central Andes Culture Area over a period of ∼1000 years (AD ∼ 50-1300) involved processes affected by population growth, warfare and hydroclimate conditions. Ortloff and Kolata [22] proposed that vulnerability to drought in pre-Columbian societies varied according to the degree of dependence on rainfed agricultural production, which is a function of environmental constraints and the development of specialized irrigation technologies to buffer drought impacts [24-26].

Compared to the Middle Horizon northern high-Andes polities (Wari and Tiwanuku centers) that deployed complex farming irrigated raised field systems across semiarid areas, complex societies of the northern sociocultural centers area in the arid Peruvian coast are expected to be more vulnerable, as farming production systems rely heavily on the local precipitation regime, subject to cycles of drought, floods and sand dune erosion [28]. We effectively verified differentiated vulnerabilities to climate risk between the northern and southern Andean sociocultural centers areas. Our results, however, contribute additional elements for understanding such a relationship between climate and societal collapse. First, hydroclimate conditions (drought and flood) were indeed a key factor in driving the growth and demise of the Moche polities, but this dynamic was greatly modulated by population size. Second, although, the long-term variations in water availability acted as an exogenous factor in the rise and collapse of the southern highland centers, it marginally affected the feedback between population growth and conflict.

Contrasting positions exist on the impact of ENSO-driven positive hydroclimate anomalies that could have affect the trajectory of past societies from the northern coast of Peru, ranging from devasting to beneficial consequences [26, 27, 29, 32, 59, 60]. Demographic dynamic models attest to a positive effect of these events on population dynamics, which is consistent with evidence for adaptive practices used to cope with the interrelated impacts of amplified water availability and flood risk. The Moche deployed distinct cultivation areas along the arid coastal escarpment, where slopes could easily accumulate rainfall runoff and then be managed through small-scale and low-investment irrigation infrastructure that was rapidly restored after major flooding events [26]. This opportunistic production regime was complemented by seizing flooding water from rivers through networks of redundant/intermittent irrigation channels [26] or transient farming in peripheral areas [27, 59]. High lithic concentrations in the SO147-106KL core (>35%, Fig. 1E) indicate that the period between AD 350 and 590 was a period favorable for thriving opportunistic and intensified maize production. This implies that positive population growth rates (AD 400-600) were achieved by increased agricultural land carrying capacity insofar as water resources became abundant, but vulnerability to flooding and sand dune deposition was also mitigated.

Negative rainfall anomalies greatly challenged coastal farming systems of the Central Andes Culture Area [26, 28, 59]. Aside from depleted water availability, the loss in arable land and crop productivity was exacerbated by desertification, flood as a consequence of strong local rainfall link to El Niño events, and localized dune transgression [59]. Infrastructure against sand encroachment, redundant irrigation canals, and high mobility were recurrent mitigation strategies used in the Moche period to face drought impacts [26, 59]. Data from the SO147-106KL core and independent multiproxy reconstructions [61-64] indicate that the Eastern Pacific entered the La Niña mean state between AD 900 and 1100, which resulted in a multicentennial drought (Fig. 1H). The demographic collapse detected in AD 900 matches closely with the chronology of the most severe drought event in the Peruvian coast of the last 4000 years [21]. The population decline from AD 650 onward (Fig. 1B, D), however, suggests that such demographic decline was felt earlier and regardless of adaptive strategies. Thus, the impact of a long-term depletion in water availability and/or arable land since AD 650 appears to be modulated by population size. This nonadditive effect through the per capita resource share [4, 5, 53, 55] implies that even minor hydroclimate changes are capable of triggering disproportionate demographic responses when population sizes and production systems are near the maximum.

Increased intensity of warfare and political fragmentation during the so-called Late Moche period (AD 700-900) have been linked to different mechanisms, such as prevailing adverse environmental conditions along the northern Peruvian coast [28, 29, 65]. The models developed here, however, suggest that neither hydroclimate, population pressure nor growth explain the dynamics of conflict incidence in the northern centers (Table S1). In this vein, our results concur with the empirical-theoretical analysis conducted by [66] on the trajectory of warfare in the Moche valley. Billman [66] concluded that the emergence of conflict was little related to the feedback between population pressure and land productivity and was instead caused by expansion toward coastal valleys of polities from the Andean highlands. Data derived from nonmetrical dental traits suggest that the influx of highland peoples into the Jequetepeque Valley occurred during the late Moche period [65, 67]. Such a proxy for genetic distance concurs with cultural data (i.e., changes in ceramic styles or burial patterns) in showing the existence of highland migrants over the valley. Ancient DNA analyses on the bioanthropological record suggest that Moche elite pertained to a particular “lowland” lineage [68], but that highlands and coastal populations maintained certain gene flux at least till the Spaniard colonization [69]. This implies that potential affinities between local and migrants in the Jequetepeque Valley (as well as in other Mochica territories) must be confirmed with further evidence such as ancient DNA. By this means, the role of external and internal tensions in fueling conflicts in the Moche tradition must remain speculative until additional information becomes available.

The spatially differentiated pattern in the mutual coupling between population growth and conflict detected here supports the notion that warfare was a localized process across the Central Andes Centers Area, and it was particularly relevant in the southern highland centers (e.g., [30, 66]. Conflict dynamics in this area have been attributed to the disintegration of sociopolitical networks [70], but more recently to increased population pressure on food production [35]. Alternatively, the increase in warfare intensity has been linked to the impact of a multicentennial drought that hit the Andean highlands between AD 900 and AD 1200 [e.g., 30, 31]. Arkush and Tung [30] indicate that weakened political infrastructure likely limited the drought mitigation strategies of high Andean communities (e.g., trade networks), in turn stoking conflicts over cropland, herds, and stores. The concurrent evidence for a prolonged drought [19, 23], infrastructure desecration [70] and sociopolitical dissolution [71] has led to the proposition that the Tiwanuku polities collapse and their further transformation was shaped by environmental crises that weakened the technological food production system (raised fields failure due to rainfall decrease), and the social structure of the state like polities, all of which fueled social instability, warfare, food supply disruption [22, 24]. Our modeling approach, however, suggests that the Tiwanaku-Wari rise and collapse involved an endogenous dynamic between population and warfare as part of coupled oscillations between these two variables. Under this dynamic, demographic growth or decline emerges as mutual feedback between population levels and internal warfare intensity in that the hydroclimate is an exogenous factor acting on the ratio of population to resource availability. This feedback loop is consistent with structural-demographic models proposed to explain growth-collapse cycles occurring in complex societies [15-17]. In this sense, our results partially agree with the causal mechanism evoked by McCool et al. [35] to explain the positive relationship between population growth and hydroclimate, and how this process is translated into the escalated lethal violence observed in the Nasca region after the Wari collapse at ∼AD 1000. Even when we attest for a positive effect of population pressure on the conflict dynamic, our results point to reciprocal influences between demographic levels and warfare intensity. That is, conflict negatively affect the population growth rates and demographic levels affects positively warfare.

The relationships between collapse, conflict and drought have been generally accepted even when there are important discrepancies among paleoclimate reconstructions on the direction and timing of high-Andean hydroclimate anomalies for over a millennium (AD 50-1300). Some reconstructions show prolonged aridity from AD 800-1200 [e.g., 19, 23]. New paleoclimate reconstructions for the Titicaca Basin [18, 20] are compatible with the direction and chronology of hydroclimate changes inferred from the SO147-106KL core [21]. Reconstructed lake levels provided by [18, 20] attest to an overall wet period from 800-1200, which was briefly interrupted by a moderate drought dated between AD 1120 and AD 1270.

In sum, the results of our study, together with recent paleoclimate data, challenge the traditional view of the links between hydroclimate conditions, conflict and population dynamics. For instance, population growth expansion (AD 600-900, Fig. 1B-C) appears to be associated with low warfare intensity and highly variable hydroclimate conditions [18, 20]. The demographic collapse (AD 900-1200), however, is not synchronous with the decadal-scale desiccation of the Andean highlands. Indeed, prevailing humid conditions to AD 1120 [18] preclude the role of environmental stress as an explanatory factor involved in the incipient increase in warfare intensity and the transition from the positive to negative population growth phase.

An important element to highlight is the theoretical models’ excellent predictive power in capturing the dynamic relationships between population, warfare, and climate change. Nevertheless, we are aware of the potential biases and sampling errors of archeological data. Our statistical models are based on indirect proxies of human population size, crop productivity, climate change and warfare intensity. Thus, these results are statistical hypotheses, and they are far from a definitive explanation. Despite these caveats, we suggest that the analysis of several chronologically affiliated datasets provides some novel insights into the patterns of warfare in parts of the Andes.

For decades, scholars and scientists have tried to understand why complex societies get to and collapse, and transform into new sociopolitical and ethnic entities [72, 73]. Our results may seem to support the view that societal changes in the Central Andes Centers are related to mutual feedback between population growth and social complexity [e.g., 74]. Although a postindustrial society has notable technological and informational advantages over ancient societies in responding to hydroclimate, the magnitude of the current population size, the consumption of energy and resources, and feedback from the climate system on a global scale should alert us to take a closer look at the social and conflict dynamics suffered by ancient societies in the face of population pressure and profound changes in climate.

## 5 Acknowledgments

This study was undertaken by the PEOPLE 3000 working group of the Past Global Changes (PAGES) project, which received support from the Swiss Academy of Sciences and Chinese Academy of Sciences. We are grateful for the extensive feedback provided by Elizabeth Arkush and Tiffiny Tung, which substantially contributed to the quality of the final draft.

## 8 Author contributions

Conceptualization: ML, EMG, SAE

Methodology: ML, EMG, SAE, AG

Investigation: ML, EMG, SAE, AG

Visualization: ML, EMG, SAE

Supervision: ML, EMG, SAE, CMS

Writing—original draft: ML, EMG, SAE

Writing—review & editing: ML, EMG, SAE, CMS

## 9 Data and materials availability

All data are available in the supplementary materials.

