## Supplemental file for "1000 years of population, warfare, and climate change in pre-Columbian societies of the Central Andes"

**This PDF file includes:**

Fig. S1

Tables S1 to S2

Data S1 to S2

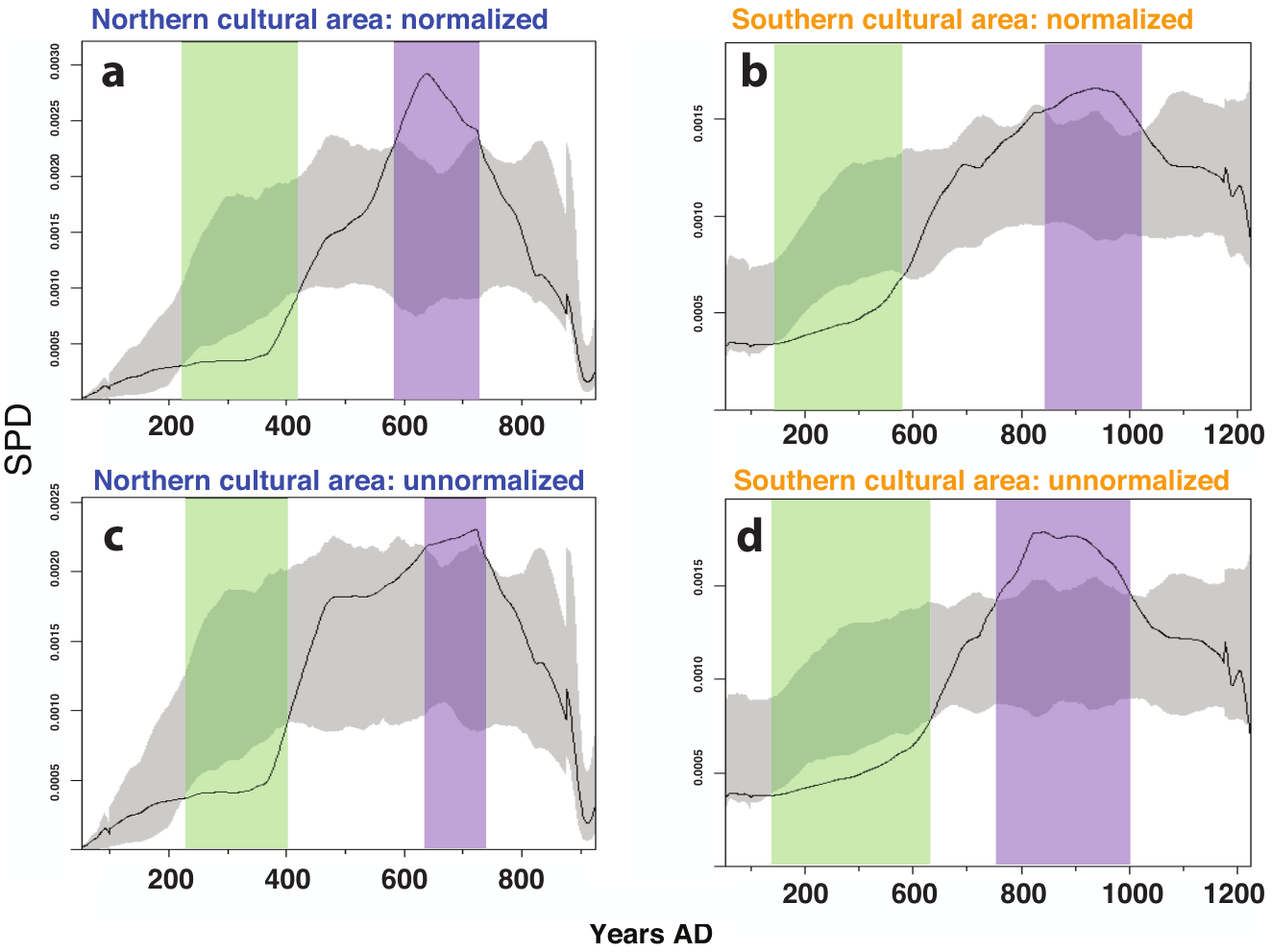

Fig. S1. Observed SPD for normalized (a-b) and unnormalized (c-d) radiocarbon dates against 95% Monte-Carlo simulated envelopes (gray areas) from fitted logistic models of population growth for the northern and southern sociocultural areas. Purple (green) vertical bars indicate positive (negative) deviations from the simulated envelope.

Table S1.: Population models (Eqs. 1-5) fitted to the time series data from the northern sociocultural area. Parameter values are given in the columns of the table. The best models were selected considering the Bayesian Akaike Information Criteria (*BIC)* and the coefficient of prediction (σ^2^ ) of the simulated total trajectory predictions initiated with the first observed value of the time series. *R^2^* is the coefficient of determination (% of explained variance) of the fitted models.

| **Population models** |  |  | **Climate** | **Warfare** |  |  |  |  |
| --- | --- | --- | --- | --- | --- | --- | --- | --- |
| Normalized | r_m_ | K | α | β | γ | BIC | R^2^ | σ^2^ |
| **Climate** | **0.25** | **-0.0016** | **0.0001^*^** |  |  | **-33.13** | **0.60** | **0.94** |
| Warfare | 0.057 | 0.0004 |  | 0.055 |  | -6.83 | 0.20 | 0.40 |
| Warfare lateral effect | 0.17 | 0.011 |  |  | -0.004 | -7.36 | 0.22 | 0.11 |
| Climate and Warfare | -1.34 | 0.004 | -0.0003^*^ | 0.65^*^ |  | -41.80 | 0.73 | 0.96 |
| Climate and Warfare | 0.25 | -0.0005 | 0.00013^*^ |  | -0.0005 | -29.73 | 0.62 | 0.95 |
| **Unnormalized** |  |  |  |  |  |  |  |  |
| **Climate** | **0.25** | **-0.0014** | **0.0001^*^** |  |  | **-33.75** | **0.60** | **0.95** |
| Warfare | -0.28 | -0.0015 |  | 0.21 |  | -12.57 | 0.27 | 0.68 |
| Warfare lateral effect | 0.21 | 0.0023 |  |  | -0.0003 | -11.77 | 0.26 | 0.57 |
| Climate and Warfare | -1.14 | 0.0026 | -0.0002^*^ | 0.58^*^ |  | -41.09 | 0.70 | 0.93 |
| Climate and Warfare | 0.29 | -0.0048 | 0.00007^*^ |  | 0.0015 | -31.82 | 0.61 | 0.95 |
| **Warfare models** |  |  |  |  |  |  |  |  |
| Normalized | μ | λ | ψ |  |  | BIC | R^2^ | σ^2^ |
| Population | -0.013 | 10.26 |  |  |  | -198.33 | 0.25 | -2.12 |
| **Population + Climate** | **-0.052** | **11.86** | **0.0011** |  |  | **-206.69** | **0.46** | **0.44** |
| Population/Climate | -0.009 | 201.88 |  |  |  | -191.68 | 0.09 | -0.67 |
| **Unnormalized** |  |  |  |  |  |  |  |  |
| Population | -0.015 | 12.49 |  |  |  | -199.80 | 0.28 | -0.59 |
| **Population + Climate** | **-0.059** | **15.30** | **0.0012** |  |  | **-211.92** | **0.53** | **0.60** |
| Population/Climate | -0.015 | 318.74 |  |  |  | -190.78 | 0.07 | -0.72 |

Table S2.

**Population models (Eqs. 1-5) fitted to the time series data from the southern sociocultural area**. Parameter values are given in the columns of the table. The best models were selected considering the Bayesian Akaike Information Criteria (*BIC)* and the coefficient of prediction (σ^2^) of the simulated total trajectory predictions initiated with the first observed value of the time series. *R^2^* is the coefficient of determination (% of explained variance) of the fitted models.

| **Population models** |  |  | **Climate** | **Warfare** |  |  |  |  |
| --- | --- | --- | --- | --- | --- | --- | --- | --- |
| Normalized | r_m_ | K | α | β | γ | BIC | R^2^ | σ^2^ |
| Climate | 0.075 | -0.0018 | 0.00014 |  |  | -85.76 | 0.33 | 0.67 |
| **Warfare** | **0.16** | **0.006** |  | **-0.078^*^** |  | **-109.63** | **0.63** | **0.87** |
| **Warfare lateral effect** | **0.087** | **0.003** |  |  | **-0.0008^*^** | **-108.54** | **0.62** | **0.75** |
| Climate and Warfare | 0.16 | 0.004 | 0.00008 | -0.076^*^ |  | -106.04 | 0.63 | 0.90 |
| Climate and Warfare | 0.098 | 0.003 | -0.00003 |  | -0.00075^*^ | -106.75 | 0.64 | 0.85 |
| **Unnormalized** |  |  |  |  |  |  |  |  |
| Climate | 0.066 | -0.0022 | 0.00016 |  |  | -76.07 | 0.30 | 0.52 |
| **Warfare** | **0.16** | **0.011** |  | **-0.091^*^** |  | **-103.72** | **0.65** | **0.87** |
| **Warfare lateral effect** | **0.075** | **0.003** |  |  | **-0.0009^*^** | **-104.81** | **0.66** | **0.69** |
| Climate and Warfare | 0.16 | 0.0006 | 0.0004 | -0.087^*^ |  | -100.44 | 0.65 | 0.90 |
| Climate and Warfare | 0.087 | 0.0032 | -0.00002 |  | -0.0008^*^ | -102.26 | 0.67 | 0.73 |
| **Warfare models** |  |  |  |  |  |  |  |  |
| Normalized | μ | λ | ψ |  |  | BIC | R^2^ | σ^2^ |
| Population | -0.059 | 99.97^*^ |  |  |  | -199.94 | 0.84 | 0.96 |
| Population + Climate | 0.002 | 75.81^*^ | -0.0015^*^ |  |  | -205.62 | 0.88 | 0.99 |
| **Population/Climate** | **-0.038** | **1699^*^** |  |  |  | **-230.29** | **0.93** | **0.99** |
| **Unnormalized** |  |  |  |  |  |  |  |  |
| Population | -0.055 | 95.08^*^ |  |  |  | -199.84 | 0.84 | 0.92 |
| Population + Climate | 0.011 | 70.66^*^ | -0.0016^*^ |  |  | -209.06 | 0.89 | 0.98 |
| **Population/Climate** | **-0.037** | **1663^*^** |  |  |  | **-236.72** | **0.94** | **0.99** |

Data S1. (separate file)

Time series data used for fitting and simulating the population dynamic models (Eqs. 1-5) from the northern sociocultural area. The columns are: the calibrated years before present (calBP), the calendar years (acdc), the normalized and non-normalized Summed Probability Distributions (SPDn and SPDnn), the respective per capita growth rates (Rn and Rnn), the lithic concentrations (Lith%), the warfare intensity (War) and the warfare growth rates (Rwar)..

| **calBP** | **acdc** | **SPDn** | **Rn** | **SPDnn** | **Rnn** | **Lith%** | **War** | **Rwar** |
| --- | --- | --- | --- | --- | --- | --- | --- | --- |
| 1950 | -1 | 0.00003 | 0.08761 | 0.00004 | 0.09149 | 38.688 | 2.9374 | -0.0306 |
| 1925 | 25 | 0.00003 | 0.35980 | 0.00004 | 0.36901 | 37.504 | 2.8488 | -0.0315 |
| 1900 | 50 | 0.00005 | 0.25903 | 0.00006 | 0.26496 | 36.938 | 2.7603 | -0.0310 |
| 1875 | 75 | 0.00006 | 0.58579 | 0.00007 | 0.59020 | 37.246 | 2.6761 | -0.0293 |
| 1850 | 100 | 0.00011 | 0.37179 | 0.00013 | 0.37320 | 38.343 | 2.5987 | -0.0268 |
| 1825 | 125 | 0.00016 | 0.15332 | 0.00019 | 0.15422 | 39.738 | 2.5300 | -0.0236 |
| 1800 | 150 | 0.00018 | 0.21538 | 0.00022 | 0.21383 | 40.655 | 2.4709 | -0.0200 |
| 1775 | 175 | 0.00023 | 0.07319 | 0.00028 | 0.06801 | 40.497 | 2.4219 | -0.0162 |
| 1750 | 200 | 0.00025 | 0.05837 | 0.00030 | 0.04891 | 39.185 | 2.3829 | -0.0123 |
| 1725 | 225 | 0.00026 | 0.09583 | 0.00031 | 0.08814 | 37.095 | 2.3537 | -0.0085 |
| 1700 | 250 | 0.00029 | 0.02956 | 0.00034 | 0.02403 | 34.874 | 2.3337 | -0.0048 |
| 1675 | 275 | 0.00030 | 0.00521 | 0.00035 | 0.00046 | 33.255 | 2.3226 | -0.0012 |
| 1650 | 300 | 0.00030 | 0.01581 | 0.00035 | 0.01344 | 32.760 | 2.3197 | 0.0021 |
| 1625 | 325 | 0.00030 | 0.09548 | 0.00035 | 0.09266 | 33.474 | 2.3247 | 0.0053 |
| 1600 | 350 | 0.00033 | 0.22545 | 0.00039 | 0.23126 | 35.116 | 2.3371 | 0.0082 |
| 1575 | 375 | 0.00042 | 0.44136 | 0.00049 | 0.45920 | 37.108 | 2.3563 | 0.0109 |
| 1550 | 400 | 0.00065 | 0.30295 | 0.00077 | 0.31245 | 38.772 | 2.3822 | 0.0133 |
| 1525 | 425 | 0.00088 | 0.22522 | 0.00106 | 0.22825 | 39.649 | 2.4140 | 0.0153 |
| 1500 | 450 | 0.00110 | 0.13828 | 0.00133 | 0.13942 | 39.758 | 2.4512 | 0.0168 |
| 1475 | 475 | 0.00126 | 0.04671 | 0.00153 | 0.00671 | 39.331 | 2.4928 | 0.0178 |
| 1450 | 500 | 0.00132 | 0.06319 | 0.00154 | 0.00224 | 38.427 | 2.5377 | 0.0181 |
| 1425 | 525 | 0.00141 | 0.11821 | 0.00154 | 0.01356 | 37.217 | 2.5841 | 0.0175 |
| 1400 | 550 | 0.00159 | 0.17652 | 0.00156 | 0.04230 | 36.008 | 2.6297 | 0.0161 |
| 1375 | 575 | 0.00189 | 0.14795 | 0.00163 | 0.05050 | 35.110 | 2.6723 | 0.0141 |
| 1350 | 600 | 0.00220 | 0.12696 | 0.00171 | 0.07221 | 34.715 | 2.7101 | 0.0117 |
| 1325 | 625 | 0.00249 | 0.02359 | 0.00184 | 0.07014 | 34.808 | 2.7419 | 0.0090 |
| 1300 | 650 | 0.00255 | -0.03668 | 0.00197 | 0.04543 | 35.021 | 2.7667 | 0.0062 |
| 1275 | 675 | 0.00246 | -0.04622 | 0.00207 | 0.03991 | 34.676 | 2.7839 | 0.0033 |
| 1250 | 700 | 0.00235 | -0.04082 | 0.00215 | 0.01493 | 33.205 | 2.7931 | 0.0003 |
| 1225 | 725 | 0.00226 | -0.14092 | 0.00218 | -0.1127 | 30.728 | 2.7940 | -0.0027 |
| 1200 | 750 | 0.00196 | -0.12166 | 0.00195 | -0.0988 | 28.051 | 2.7864 | -0.0058 |
| 1175 | 775 | 0.00173 | -0.14070 | 0.00177 | -0.0989 | 25.828 | 2.7704 | -0.0089 |
| 1150 | 800 | 0.00151 | -0.23539 | 0.00160 | -0.1376 | 24.205 | 2.7459 | -0.0119 |
| 1125 | 825 | 0.00119 | -0.08201 | 0.00139 | -0.0821 | 22.865 | 2.7134 | -0.0149 |
| 1100 | 850 | 0.00110 | -0.25679 | 0.00128 | -0.2552 | 21.488 | 2.6733 | -0.0176 |
| 1075 | 875 | 0.00085 | -0.28293 | 0.00100 | -0.2792 | 20.145 | 2.6266 | -0.0200 |
| 1050 | 900 | 0.00064 | -0.34403 | 0.00075 | -0.3479 | 19.096 | 2.5746 | -0.0217 |
| 1025 | 925 | 0.00045 | -0.34461 | 0.00053 | -0.3526 | 18.500 | 2.5194 | -0.0224 |

Data S2. Time series data used for fitting and simulating the population dynamic models (Eqs. 1-5) from the southern sociocultural area. The columns are: the calibrated years before present (calBP), the calendar years (acdc), the normalized and non-normalized Summed Probability Distributions (SPDn and SPDnn), the respective per capita growth rates (Rn and Rnn), the lithic concentrations (Lith%), the warfare intensity (War) and the warfare growth rates (Rwar).

| calBP | acdc | SPDn | Rn | SPDnn | Rnn | Lith% | War | Rwar |
| --- | --- | --- | --- | --- | --- | --- | --- | --- |
| 1650 | 300 | 0.0003 | 0.0095 | 0.00034 | 0.00691 | 35.471 | 1.0686 | -0.0329 |
| 1625 | 325 | 0.0003 | 0.0170 | 0.00035 | 0.01118 | 35.704 | 1.0339 | -0.0272 |
| 1600 | 350 | 0.0003 | 0.0429 | 0.00035 | 0.03404 | 36.245 | 1.0062 | -0.0219 |
| 1575 | 375 | 0.0003 | 0.0470 | 0.00036 | 0.03283 | 36.929 | 0.9844 | -0.0175 |
| 1550 | 400 | 0.0003 | 0.0358 | 0.00038 | 0.02419 | 37.562 | 0.9673 | -0.0139 |
| 1525 | 425 | 0.0004 | 0.0388 | 0.00038 | 0.02793 | 37.991 | 0.9540 | -0.0111 |
| 1500 | 450 | 0.0004 | 0.0299 | 0.00040 | 0.02185 | 38.169 | 0.9435 | -0.0091 |
| 1475 | 475 | 0.0004 | 0.0574 | 0.00040 | 0.04086 | 38.110 | 0.9350 | -0.0077 |
| 1450 | 500 | 0.0004 | 0.0789 | 0.00042 | 0.05545 | 37.819 | 0.9277 | -0.0068 |
| 1425 | 525 | 0.0004 | 0.1153 | 0.00044 | 0.06628 | 37.334 | 0.9214 | -0.0062 |
| 1400 | 550 | 0.0005 | 0.1692 | 0.00048 | 0.08748 | 36.731 | 0.9157 | -0.0056 |
| 1375 | 575 | 0.0006 | 0.1656 | 0.00052 | 0.08431 | 36.081 | 0.9107 | -0.0046 |
| 1350 | 600 | 0.0007 | 0.2151 | 0.00056 | 0.15925 | 35.428 | 0.9064 | -0.0032 |
| 1325 | 625 | 0.0008 | 0.1372 | 0.00066 | 0.19683 | 34.760 | 0.9036 | -0.0008 |
| 1300 | 650 | 0.0010 | 0.0852 | 0.00081 | 0.16002 | 33.995 | 0.9029 | 0.0027 |
| 1275 | 675 | 0.0011 | 0.0491 | 0.00095 | 0.11650 | 32.989 | 0.9054 | 0.0077 |
| 1250 | 700 | 0.0011 | -0.0090 | 0.00106 | 0.04189 | 31.633 | 0.9124 | 0.0143 |
| 1225 | 725 | 0.0011 | 0.0568 | 0.00111 | 0.10305 | 29.972 | 0.9255 | 0.0226 |
| 1200 | 750 | 0.0012 | 0.0435 | 0.00123 | 0.07382 | 28.197 | 0.9467 | 0.0325 |
| 1175 | 775 | 0.0012 | 0.0373 | 0.00132 | 0.07003 | 26.483 | 0.9780 | 0.0437 |
| 1150 | 800 | 0.0013 | 0.0437 | 0.00142 | 0.08851 | 24.907 | 1.0216 | 0.0555 |
| 1125 | 825 | 0.0013 | 0.0088 | 0.00155 | -0.00364 | 23.450 | 1.0800 | 0.0673 |
| 1100 | 850 | 0.0013 | 0.0255 | 0.00154 | -0.01572 | 22.084 | 1.1552 | 0.0783 |
| 1075 | 875 | 0.0014 | 0.0224 | 0.00152 | -0.00045 | 20.845 | 1.2492 | 0.0877 |
| 1050 | 900 | 0.0014 | 0.0119 | 0.00152 | -0.02158 | 19.799 | 1.3637 | 0.0948 |
| 1025 | 925 | 0.0014 | 0.0004 | 0.00149 | -0.04304 | 18.982 | 1.4993 | 0.0993 |
| 1000 | 950 | 0.0014 | -0.0138 | 0.00142 | -0.04194 | 18.375 | 1.6558 | 0.1008 |
| 975 | 975 | 0.0014 | -0.0613 | 0.00137 | -0.09166 | 17.926 | 1.8314 | 0.0992 |
| 950 | 1000 | 0.0013 | -0.0795 | 0.00125 | -0.09642 | 17.594 | 2.0223 | 0.0943 |
| 925 | 1025 | 0.0012 | -0.0798 | 0.00113 | -0.07081 | 17.359 | 2.2223 | 0.0863 |
| 900 | 1050 | 0.0011 | -0.0756 | 0.00105 | -0.04461 | 17.215 | 2.4226 | 0.0760 |
| 875 | 1075 | 0.0010 | -0.0112 | 0.00101 | -0.00441 | 17.179 | 2.6138 | 0.0650 |
| 850 | 1100 | 0.0010 | -0.0002 | 0.00100 | -0.00680 | 17.277 | 2.7892 | 0.0540 |
| 825 | 1125 | 0.0010 | -0.0243 | 0.00100 | -0.03742 | 17.535 | 2.9441 | 0.0436 |
| 800 | 1150 | 0.0010 | -0.0517 | 0.00096 | -0.07365 | 18.007 | 3.0754 | 0.0338 |
| 775 | 1175 | 0.0010 | -0.1246 | 0.00089 | -0.16630 | 18.733 | 3.1812 | 0.0246 |
| 750 | 1200 | 0.0008 | -0.1322 | 0.00076 | -0.18269 | 19.688 | 3.2604 | 0.0158 |
| 725 | 1225 | 0.0007 | -0.2375 | 0.00063 | -0.28018 | 20.761 | 3.3121 | 0.0072 |
| 700 | 1250 | 0.0006 | -0.2819 | 0.00048 | -0.32134 | 21.826 | 3.3361 | -0.0013 |
